# The expression and function of 28s5-rtsRNA: an update

**DOI:** 10.1101/2022.05.29.492757

**Authors:** Yi Wang, Yanjie Zhao, Shuai Li

## Abstract

Our previous research identified an abundant 28s rRNA 5’ terminal derived small RNA (named 28s5-rtsRNA), which could decrease multiple ribosomal protein mRNA levels. In the last several years, the development of novel high-throughput RNA sequencing methods, exampled as PANDORA-seq, has changed the repertoire of small RNAs. In the present study, we reanalyzed the PANDORA-seq data by using a sRNAbench bioinformatic tool and revealed that 28s5-rtsRNA was among the most abundant small RNAs, particularly after T4PNK treatment during sequencing library preparation. 28s5-rtsRNA pairing with 5.8s rRNA 3’ terminal derived small RNA could also decrease multiple ribosomal mRNA levels. At last, we performed a mutational scan to identify the key nucleotides involved in the 28s5-rtsRNA duplex function.

## Introduction

Along with the development of small RNA high-throughput sequencing (HTS) technology, the repertoire of small RNAs was greatly expanded [1-5]. In addition to well-characterized microRNAs and piRNAs, small RNAs were found to be originated from abundant housekeeping non-coding RNAs, such as tRNAs, rRNAs, and snoRNAs [1-6]. Recently, ample research has been focused on tRNA-derived small RNAs (tsRNAs) and revealed its diverse functions, including promoting stress-induced translational repression [7, 8], mediating epigenetic inheritance of paternally acquired traits [9-11], inhibiting cell apoptosis [12] and so on [13, 14]. Meanwhile, multiple studies have identified rRNA-derived small RNAs (rsRNAs) by HTS [2, 15-17]. For example, the expression of rsRNAs was reported to be associated with DNA damage and inflammatory responses [16, 18].

Our previous research identified 28s rRNA 5’ terminal derived small RNA (named 28s5-rtsRNA) as the most abundant rRNA-derived small RNAs [19]. We found that 28s5-rtsRNA overexpression could decrease multiple ribosomal protein mRNA levels and alter the 28s/18s rRNA ratio. Here by using a sRNAbench bioinformatic tool, we analyzed PANDORA-seq data, which overcomes RNA modifications during sequencing library construction, and demonstrated that 28s5-rtsRNA might be the most abundant small RNA in the whole small RNA repertoire. We also revealed that 28s5-rtsRNA could perform its function through paring with other small RNAs, e.g., 5.8s rRNA 3’ terminal derived small RNA (5.8s3-rtsRNA). At last, a mutational scan was performed to identify key nucleotides involved in 28s5-rtsRNA function.

## Methods and Materials

### Cell culture

The HeLa cell line was maintained in DMEM cell culture medium (Thermo Fisher Scientific, Hudson, NH, USA), supplemented with 10% FBS and 1% Penicillin-Streptomycin Solution at 37 □ with CO2.

### Small RNA high throughput sequencing data analysis

sRNAbench bioinformatic tool was employed to analyze the small RNA high throughput sequencing data using default parameters [20]. The raw SRA files were subjected to analysis. The high throughput sequencing data used in this study were listed in Supplementary Table 1. After adapter trimming, length and quality filtering and collapsing, a *reads_orig*.*fa* file, containing small RNA sequence and read count, was generated for downstream analysis. 28s5-rtsRNA and 28s5-rtsRNA half were identified by in-house Perl scripts.

### RNA transfection

The RNA sequences used in this study are listed in Supplementary Table 2. The duplex 28s5-5.8s3-rtsRNA and mutational isoforms were annealed to form duplexes. Single-stranded small RNA and duplex RNA were transfected using RNAiMax (Thermo Fisher Scientific) and Lipofectamine 2000 (Thermo Fisher Scientific), respectively.

### RNA-sequencing and data processing

Total RNA was extracted using RNAios Plus reagent (TaKaRa, Kusatsu, Japan), and 28s/18s rRNA ratios were analyzed by Agilent 2100 (Agilent Technologies, CA, USA). RNA-sequencing was performed as previously described [19]. SeqMonk (v1.41.0, Babraham Bioinformatics) software was employed to visualize and analyze RNA-sequencing generated BAM files. Gene Ontology (GO) analyses were completed using ToppGene.

### Real-time PCR

RPS13 mRNA levels were analyzed by real-time PCR using GAPDH mRNA as control. The real-time PCR was performed as previously described [19].

### Statistics

For RNA-sequencing data, DESeq2 was used to identify differentially expressed genes. Real-time PCR relative mRNA expression levels were presented as the mean ± s.e.m. Differences were assessed by two-tailed Student’s t-test using GraphPad software. *p* < 0.05 was considered to be statistically significant.

## Results

### 28s5-rtsRNA is one of the most abundant small RNA as revealed by PANDORA-seq

During small RNA sequencing library preparation, RNA modifications, such as 3’ terminal modifications (3’-phosphate, 2’,3’-cyclic phosphate) and RNA methylations (m^1^A, m^3^C, m^1^G), could interfere adapter ligation and reverse transcription leading to biased sequencing results [2, 4, 5]. This is particularly severe for small RNAs originated from tRNAs and rRNAs. These housekeeping non-coding RNAs undergo intensive RNA modifications during their maturation [21, 22]. The enzymatical treatments to remove these RNA modifications during library construction greatly changed the repertoires of small RNAs revealed by high-throughput sequencing [2, 3]. For example, Sharma *et al*. employed polynucleotide kinase (PNK) to resolute cyclic 2’-3’ phosphates left at the 3’ end of RNase A and RNase T family cleavage products [23]. This treatment resulted in a dramatic increase in rRNA-mapping reads. As shown in Supplementary Table 3 and 4, 28s5-rtsRNA is the most abundant small RNA after PNK treatment. The RPM (read per million reads) value of 28s5-rtsRNA increased from 26,765 to 529,563 in mouse cauda epididymis sperm small RNAs after PNK treatment.

To systematically eliminate the impact of RNA modifications on small RNA sequencing library construction, Shi *et al*. developed the PANDORA-seq method by employing a combinatorial enzymatic treatment to remove RNA modifications that block adapter ligation and reverse transcription. Here, by using a sRNAbench bioinformatic tool, we analyzed the PANDORA-seq data. As shown in Table 1, 28s5-rtsRNA serves as the most abundant small RNAs in mouse brain. Five out of the top ten small RNAs are 28s5-rtsRNAs with different lengths, ranging from 34-38bp. Similarly, 28s5-rtsRNAs are the most abundant small RNAs in mouse liver (Supplementary Table 5) and human ECS cells (Supplementary Table 6). Inconsistent with the results from Sharma *et al*., PNK treatment could increase the RPM values of 28s5-rtsRNAs in mouse brain, liver, and ESC (Figure 1). Over 90% of 28s5-rtsRNAs terminate within an AUUUA region (Figure 2AB).

**Figure 1.**
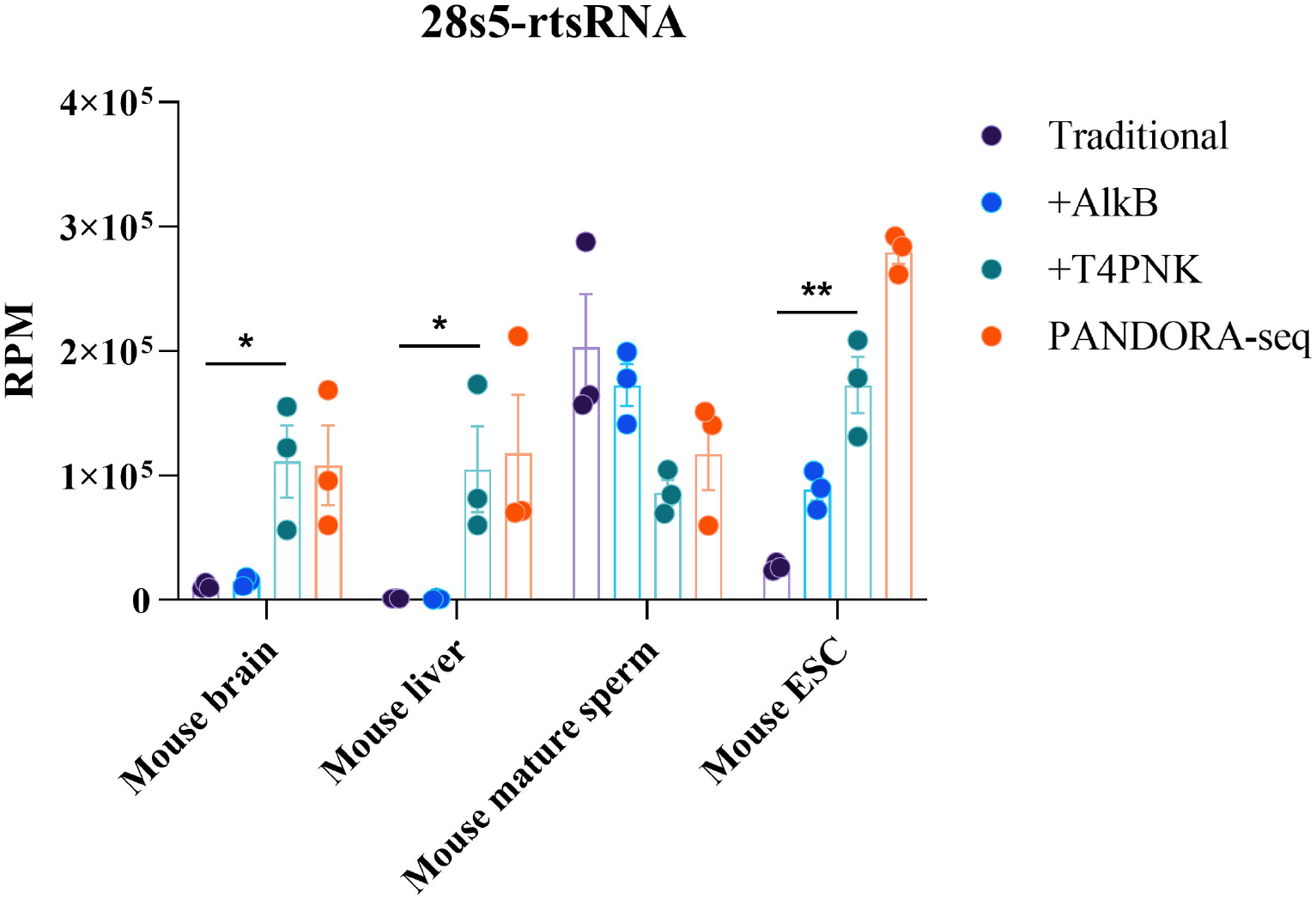
Tissue specific expression of 28s5-rtsRNA in mice under traditional RNA-seq, AlkB-facilitated RNA-seq, T4PNK-facilityed RNA-seq and PANDORA-seq. The raw small RNA sequencing data were processed by the sRNAbench bioinformatic tool. The RPM values of 28s5-rtsRNA were present. n = 3. The data represent mean ± s.e.m. \**P* < 0.05; \*\**P* < 0.01; statistical significance calculated using two-tailed Student’s t-test.

**Figure 2.**
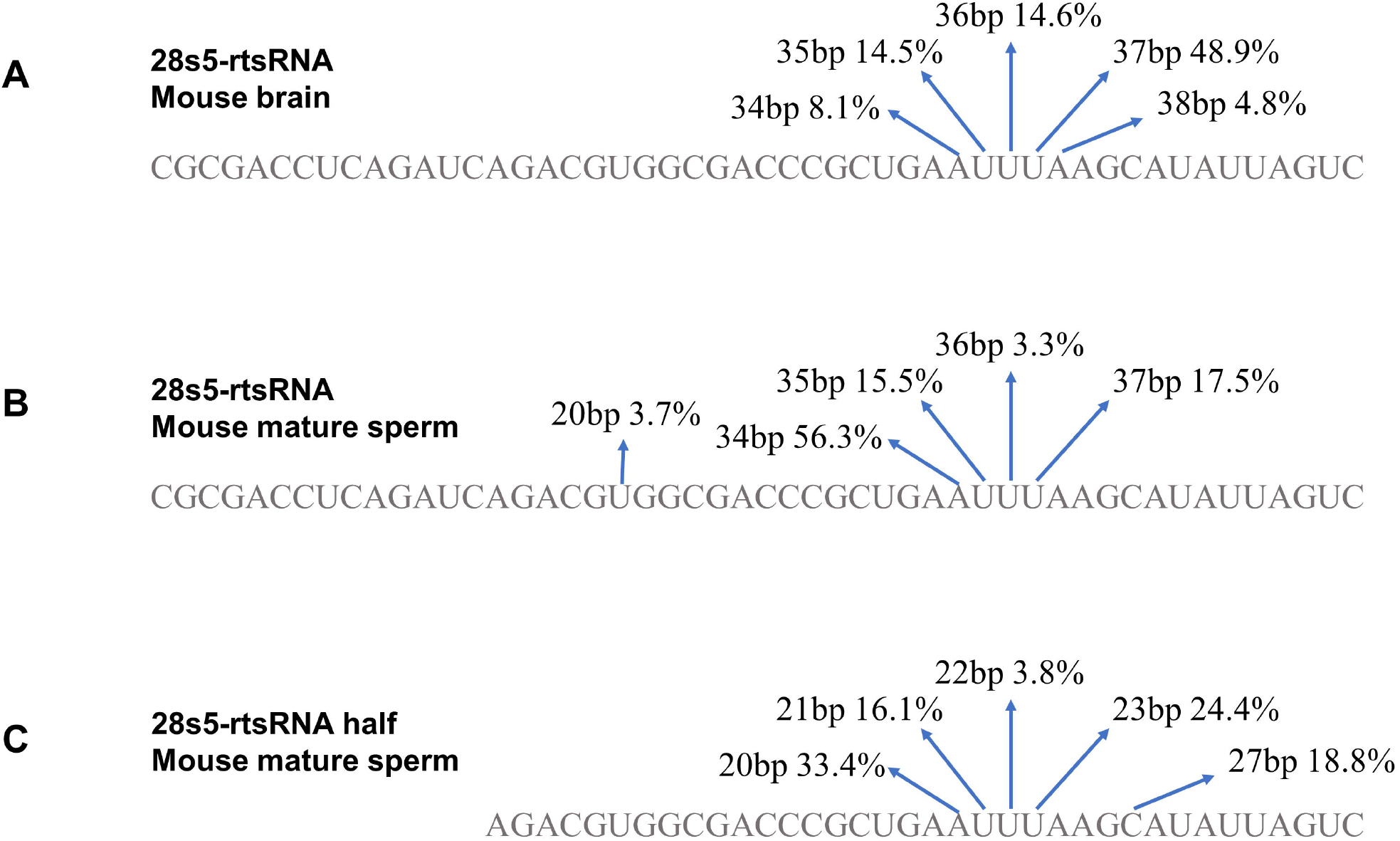
Statistics of the 3’ terminals of 28s5-rtsRNA and 28s5-rtsRNA half. The statistics of the 3’ terminal of 28s5-rtsRNA in mouse brain (A) and mouse mature sperm (B) were present. The statistics of the 3’ terminal of 28s5-rtsRNA half in mouse mature sperm (C). The percentages of specific 28s5-rtsRNAs termination were indicated by arrows.

For mouse mature sperm, three out of the top ten small RNAs are 28s5-rtsRNAs (Table 2). Moreover, four out of the top ten small RNAs are fragments originated from 28s5-rtsRNAs starting from its 15^th^ nucleotide, e.g., the second most abundant small RNA AGACGUGGCGACCCGCUGAA. We here refer to these sequences as 28s5-rtsRNA half. PNK treatment could increase the RPM values of 28s5-rtsRNA half in sperm but not that of the 28s5-rtsRNA (Figure 1, Figure 3). Over 75% of 28s5-rtsRNA half terminate within the AUUUA region (Figure 2C).

**Figure 3.**
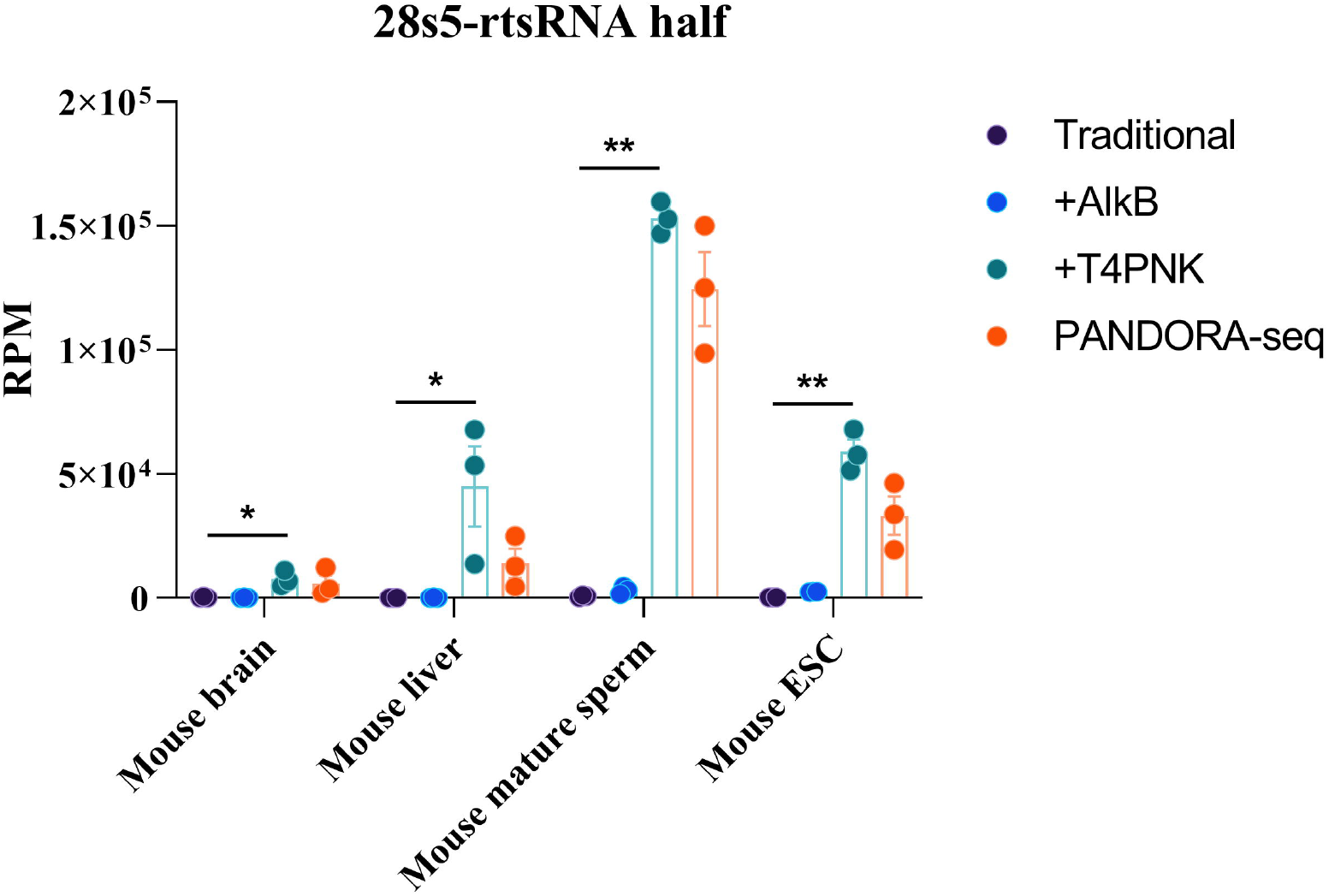
Tissue specific expression of 28s5-rtsRNA half in mice under traditional RNA-seq, AlkB-facilitated RNA-seq, T4PNK-facilityed RNA-seq and PANDORA-seq. The raw small RNA sequencing data were processed by the sRNAbench bioinformatic tool. The RPM values of 28s5-rtsRNA half were present. n = 3. The data represent mean ± s.e.m. \**P* < 0.05; \*\**P* < 0.01; statistical significance calculated using two-tailed Student’s t-test.

### 28s5-rtsRNA could inhibit ribosomal protein mRNA levels and increase 28s/18s rRNA ratios when paring with 5.8s3-rtsRNA

Our previous results demonstrated that 28s5-rtsRNA duplex (paring with a complementary single-stranded RNA, Figure 4B) could upregulate 28s/18s rRNA ratios and downregulate multiple ribosomal protein mRNA levels. During rRNA maturation processing, the 5’ terminal of 28s rRNA must base pair to the 3’ terminal of 5.8s rRNA. This duplex retains and is located on the surface of mature ribosomes (Figure 4A) [24]. In the present study, we annealed 28s5-rtsRNA with 5.8s rRNA 3’ terminal derived small RNA (refers as 5.8s3-rtsRNA) to form a novel duplex (Figure 4B). We transfected this 28s5-5.8s3-rtsRNA-duplex into the HeLa cells and found an upregulation of 28s/18s rRNA ratios compared with the Negative Control transfection group (Figure 4C-E).

**Figure 4.**
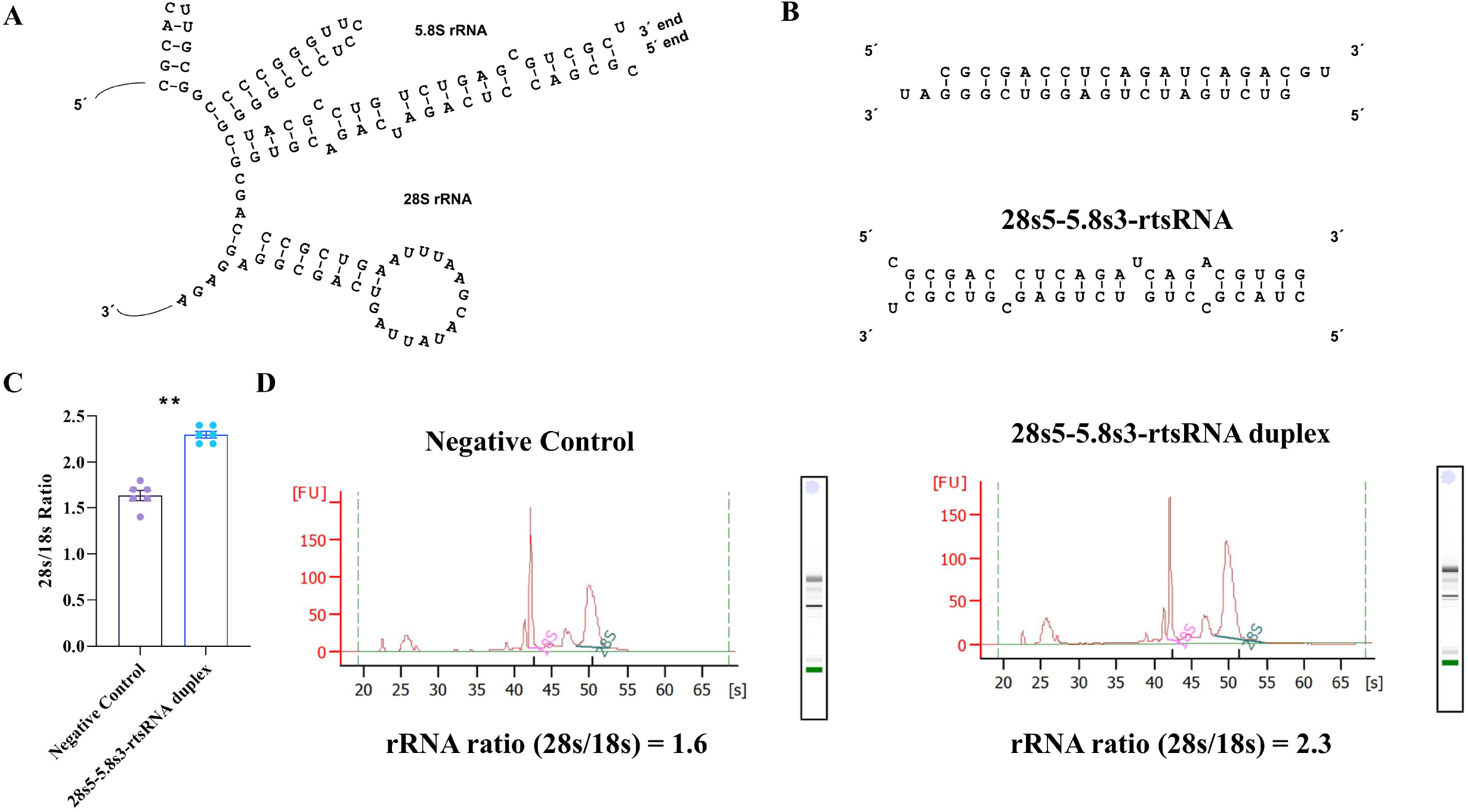
28s5-rtsRNA duplexing and impact on 28s/18s rRNA ratio. (A) Base pairing between human 28s and 5.8s rRNAs. (B) 28s5-rtsRNA base pairing with complementary and 5.8s3-rtsRNA partner single-stranded RNAs. (C) 28s/18s rRNA ratio in Negative Control and 28s5-5.8s3-rtsRNA transfected HeLa cells. n = 6. The data represent mean ± s.e.m. \*\**P* < 0.01; statistical significance calculated using two-tailed Student’s t-test. (D) Reprehensive images of RNA size distribution from Negative Control and 28s5-5.8s3-rtsRNA transfected HeLa cells analyzed by Agilent Bioanalyzer.

We then performed RNA-sequencing to explore the transcriptomic dynamics after the transfection of the 28s5-5.8s3-rtsRNA-duplex. There are 18 ribosomal protein large (RPL) and 10 ribosomal protein small (RPS) genes that are significantly downregulated (Supplementary Table 7). Inconsistent with our previous results, RPS13 has a maximum downregulation among these ribosomal protein genes (Figure 5, Supplementary Figure 1).

**Figure 5.**
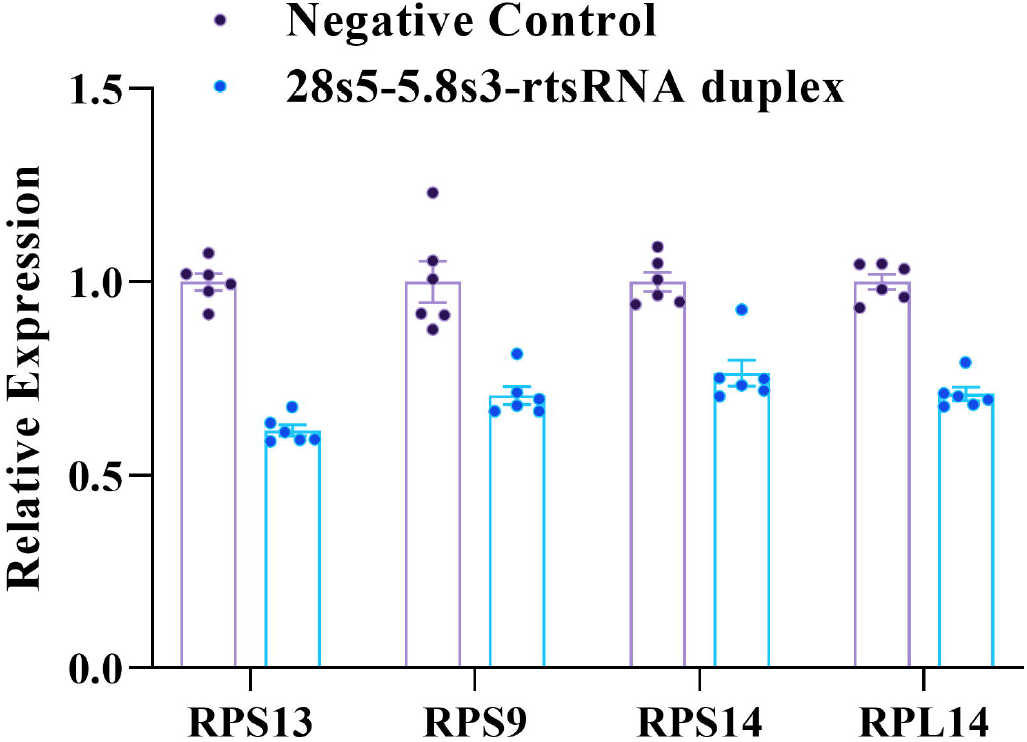
Relative gene expression by RNA-sequencing. n = 6. The data represent mean ± s.e.m.

Gene Ontology (GO) analyses on the 28s5-5.8s3-rtsRNA duplex downregulated mRNAs showed significant enrichments of cellular components associated with ribosome and lysosome membrane (Supplementary Table 8). While for the 28s5-5.8s3-rtsRNA duplex upregulated mRNAs, there are significant enrichments of the cellular component of spindle and chromosome (Supplementary Table 9). The KEGG pathway enrichment analysis revealed that the 28s5-5.8s3-rtsRNA downregulated genes were highly associated with several pathways, including lysosome, ribosome, and glutathione metabolism (Table 3). For 28s5-5.8s3-rtsRNA upregulated genes, the enriched pathways are cell cycle, RNA transport, and ribosome biogenesis (Table 4).

We also transfected the HeLa cell with single-stranded 28s5-rtsRNA. The overexpression of single-stranded 28s5-rtsRNA of different lengths could not change the 28s/18s rRNA ratios (Supplementary Figure 2A). Meanwhile, the expression levels of RPS13 mRNA were not changed by single-stranded 28s5-rtsRNA overexpression as revealed by RNA-seq (Supplementary Figure 2B).

### Mutational scan of 28s5-5.8s3-rtsRNA duplex

We next performed a mutational scan of the 28s5-5.8s-rtsRNA duplex. A total of ten mutational duplexes were independently transfected into the HeLa cell to analyze the 28s/18s rRNA ratios and RPS13 expression (Figure 6). Three terminal nucleotide deletion mutational isoforms (Mut1-Mut3) were included in the mutational scans (Figure 6A). Taking Mut1 as an example, the first 5’ terminal nucleotide C of 28s5-rtsRNA was deleted. Seven intermediate nucleotide switch mutational isoforms (Mut4-Mut10) were generated. Taking Mut4 as an example, the third nucleotide of 28s5-rtsRNA changed from C to G, and the twentieth nucleotide of 5.8s3-rtsRNA changed from G to C. This switch changed the nucleotide sequence without disrupting the duplex base paring. The sequences and base paring formats of these mutational isoforms were present in Supplementary Table 2 and Supplementary Figure 3, respectively. As shown in Figure 6B, Mut2 and Mut7-10 could increase the 28s/18s rRNA ratios. Meanwhile, as revealed by Realtime PCR and RNA-seq, Mut2 and Mut7-10 could inhibit the RPS13 mRNA levels (Figure 6CD). Thus, the first seven 5’ terminal nucleotides of 28s5-rtsRNA are important for its function.

**Figure 6.**
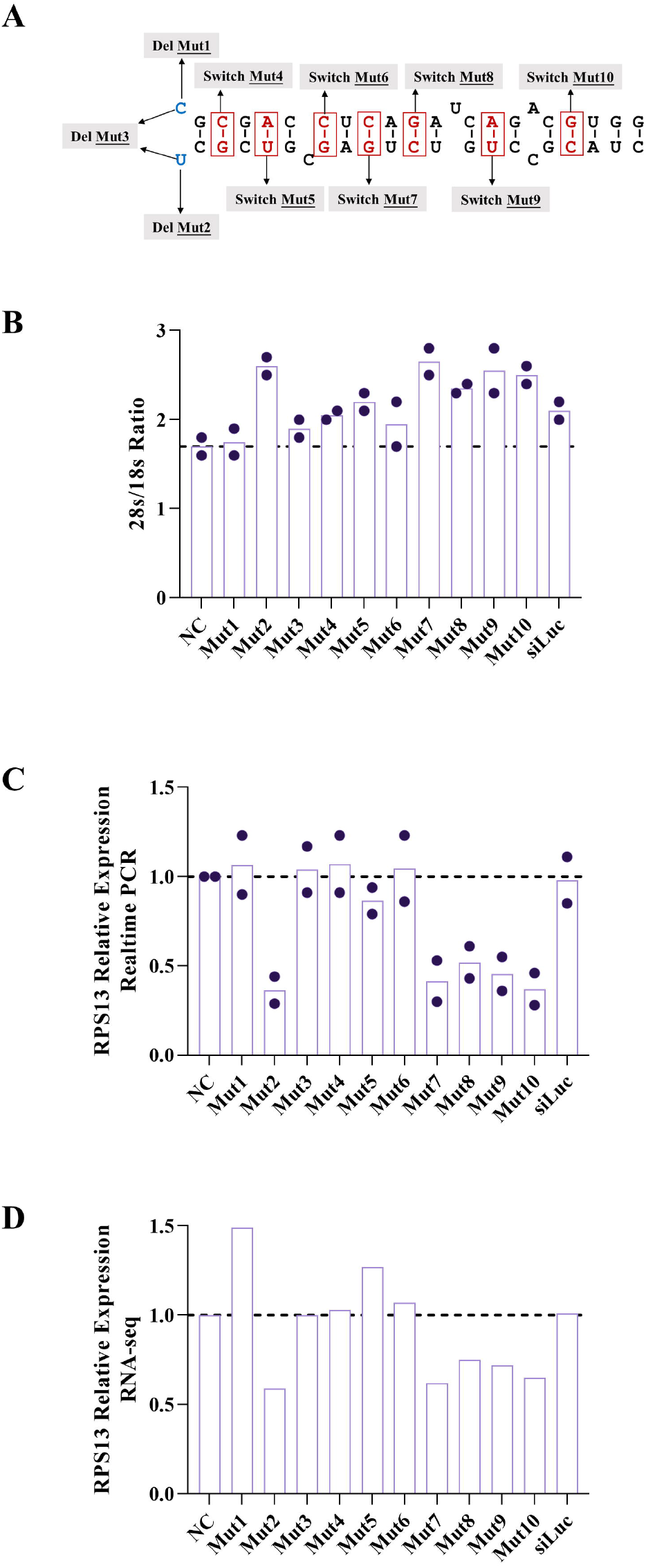
Mutational scan of 28s5-5.8s3-rtsRNA duplex. (A) Schematic diagram showing the mutations of 28s5-5.8s3-rtsRNA duplex involved in the present research. (B) 28s/18s rRNA ratios of HeLa cell transfected with different mutational duplexes. n = 2. (C) Realtime PCR analysis of RPS13 expression in HeLa cell transfected with different mutational duplexes. GAPDH mRNA levels were used for normalization. n = 2. (D) RNA-sequencing analysis of RPS13 expression. n = 1.

## Discussion

In the present study, we focused on a kind of 28s rRNA 5’ terminal derived small RNAs, referred to as 28s5-rtsRNAs. These 28s5-rtsRNAs have an identical 5’ end and different 3’ ends, leading to a length dynamic ranging from 34-38bp. The novel small RNA sequencing technology, such as PANDORA-seq, revealed that 28s5-rtsRNAs serve as one of the most abundant small RNAs. As shown in Figure 1, 28s5-rtsRNAs account for 10% - 30% of total small RNAs. The T4PNK treatments, which could resolve 2’-3’ cyclic phosphates left at the 3’ end of RNaseA and RNaseT family cleavage products, could significantly increase the RPM values of 28s5-rtsRNAs. Moreover, over 90% of 28s5-rtsRNAs terminate at the AUUUA region (Figure 2). The U and A are the preferred residues of RNaseA and RNaseT, respectively. All these information suggests RNaseA and RNaseT as the candidate enzymes involved in 28s5-rtsRNAs generation.

We also identified another high abundant 28s rRNA 5’ terminal derived small RNA (28s5-rtsRNA half), which starts at the fifteenth residue and terminates at the AUUUA region (Figure 2C). In sperm, 28s5-rtsRNA half serves as the most abundant small RNA as revealed by PANDORA-seq (Table 2). Similar with 28s5-rtsRNAs, T4PNK treatment could lead to a significant upregulation of the RPM values of 28s5-rtsRNA half (Figure 3). There is a possibility that 28s5-rtsRNA half is generated by the cleavage of 28s5-rtsRNA between its fourteen to fifteenth residues. The other half of the cleavage product should be a 14bp small RNA. However, as the deposited PANDORA-seq data filtered out small RNA sequences shorter than 15bp, we could not analyze the expression of this 14bp small RNA. Ninomiya *et al*. have demonstrated the abundant expression of 14bp 28s5-rtsRNA in the PBMC from the myeloma patients [25].

28s5-rtsRNA duplexes, formed with complementary single-stranded RNA or with 5.8s3-rtsRNA, could increase the 28s/18s rRNA ratios and downregulate multiple ribosomal protein mRNA levels (Figure 4CD). However, overexpression of single-stranded 28s5-rtsRNA could not lead to these changes (Supplementary Figure 2). Mutational scan results showed that the first seven residues of 28s5-rtsRNA are important for its function (Figure 6). However, there are several issues that need to be further addressed. First, in physiology conditions, 28s5-rtsRNA exists in single-strand form or duplex form? Second, what is 28s5-rtsRNA’s partner single-stranded RNA to form duplex? Third, what is the function of 28s5-rtsRNA?

Both 28s5-rtsRNA and 28s5-rtsRNA half are abundant in mature sperm as revealed by PANDORA-seq (Table 2). It is reported that small RNAs could be trafficked from the epididymis to developing mammalian sperm through epididymosomes thereby changing the sperm RNA payload from piRNAs to rsRNAs and tsRNAs [23]. We here analyzed the small RNA profiles of epididymosomes and found 28s5-rtsRNA served as the most abundant small RNAs (data not shown) [26]. However, the 28s5-rtsRNA half is not enriched in epididymosomes. The high abundant 28s5-rtsRNA duplex payload in sperm may help to devoid of the ribosome and strip down cytoplasm during spermatogenesis. The low temperature of testis may help to keep the duplex format of 28s5-rtsRNA. The sperm small RNAs are delivered to the zygote at fertilization, where the higher temperature resolves the duplex and mutes its duplex-associated functions (Supplementary Figure 4). Thus 28s5-rtsRNA duplex may serve as a thermometer with important regulatory functions. Moreover, sperms could swim up the temperature gradient in the oviduct [27]. This is called the sperm thermotaxis [28, 29]. 28s5-rtsRNA duplexing might be the candidate temperature sensing mechanism (Supplementary Figure 4). However, what is the partner single-stranded RNA of sperm 28s5-rtsRNA need to be further investigated. The 28s5-rtsRNA duplex may form intra-or inter-molecularly (Supplementary Figure 5).

In summary, we identified 28s5-rtsRNA as one of the most abundant small RNAs as revealed by PANDORA-seq. 28s5-5.8s3-rtsRNA duplex could increase the 28s/18s rRNA ratios and downregulate multiple ribosomal protein mRNA levels. Mutational scan experiments identified the key residues within the 28s5-5.8s3-rtsRNA duplex associated with its function.

## Acknowledgements

This work was supported by the National Natural Science Foundation of China [31870860, 31400673 to S.L.].

## Author Contributions

S.L. conceived and supervised the study. S.L. designed the experiments and analyzed the data. S.L., Y.Z., and Y.W. performed the experiments, prepared the figures, and wrote the paper. All authors discussed the results and reviewed the manuscript.

## Competing Interests

The authors have declared that no competing interest exists.

## Figure Legends

**Supplementary Figure 1**. Realtime PCR analysis of ribosomal protein mRNA levels in HeLa cells transfected with Negative Control and 28s5-5.8s3-rtsRNA duplex. GAPDH mRNA levels were used for normalization. n = 3. The data represent mean ± s.e.m. \*\**P* < 0.01; statistical significance calculated using two-tailed Student’s t-test.

**Supplementary Figure 2**. The impact of single-stranded 28s5-rtsRNAs overexpression on 28s/18s rRNA ratios (A) and RPS13 expression (B). HeLa cells were transfected with single-stranded Negative Control and 28s5-rtsRNAs with different lengths. RPS13 expression was analyzed by RNA-sequencing. n = 3. The data represent mean ± s.e.m.

**Supplementary Figure 3**. Sequences of 28s5-5.8s3-rtsRNA mutational duplexes used in this study. The switched nucleotides were marked in red in Mut4-Mut10.

**Supplementary Figure 4**. A model for 28s5-rtsRNA duplex in sperm development and thermo-sensing.

**Supplementary Figure 5**. The potential duplexing forms of 28s5-rtsRNA. The intramolecular or intermolecular duplexes of 28s5-rtsRNAs were present.

## Notes

### Competing Interest Statement

The authors have declared no competing interest.

## References

1. Shi, J., Zhou, T. & Chen, Q. (2022) Exploring the expanding universe of small RNAs, Nat Cell Biol. 24, 415–423.

2. Shi, J., Zhang, Y., Tan, D., Zhang, X., Yan, M., Zhang, Y., Franklin, R., Shahbazi, M., Mackinlay, K., Liu, S., Kuhle, B., James, E. R., Zhang, L., Qu, Y., Zhai, Q., Zhao, W., Zhao, L., Zhou, C., Gu, W., Murn, J., Guo, J., Carrell, D. T., Wang, Y., Chen, X., Cairns, B. R., Yang, X. L., Schimmel, P., Zernicka-Goetz, M., Cheloufi, S., Zhang, Y., Zhou, T. & Chen, Q. (2021) PANDORA-seq expands the repertoire of regulatory small RNAs by overcoming RNA modifications, Nat Cell Biol. 23, 424–436.

3. Cozen, A. E., Quartley, E., Holmes, A. D., Hrabeta-Robinson, E., Phizicky, E. M. & Lowe, T. M. (2015) ARM-seq: AlkB-facilitated RNA methylation sequencing reveals a complex landscape of modified tRNA fragments, Nat Methods. 12, 879–84.

4. Zheng, G., Qin, Y., Clark, W. C., Dai, Q., Yi, C., He, C., Lambowitz, A. M. & Pan, T. (2015) Efficient and quantitative high-throughput tRNA sequencing, Nat Methods. 12, 835–837.

5. Wang, H., Huang, R., Li, L., Zhu, J., Li, Z., Peng, C., Zhuang, X., Lin, H., Shi, S. & Huang, P. (2021) CPA-seq reveals small ncRNAs with methylated nucleosides and diverse termini, Cell Discov. 7, 25.

6. Ender, C., Krek, A., Friedlander, M. R., Beitzinger, M., Weinmann, L., Chen, W., Pfeffer, S., Rajewsky, N. & Meister, G. (2008) A human snoRNA with microRNA-like functions, Mol Cell. 32, 519–28.

7. Ivanov, P., Emara, M. M., Villen, J., Gygi, S. P. & Anderson, P. (2011) Angiogenin-induced tRNA fragments inhibit translation initiation, Mol Cell. 43, 613–23.

8. Fu, H., Feng, J., Liu, Q., Sun, F., Tie, Y., Zhu, J., Xing, R., Sun, Z. & Zheng, X. (2009) Stress induces tRNA cleavage by angiogenin in mammalian cells, FEBS Lett. 583, 437–42.

9. Chen, Q., Yan, M., Cao, Z., Li, X., Zhang, Y., Shi, J., Feng, G. H., Peng, H., Zhang, X., Qian, J., Duan, E., Zhai, Q. & Zhou, Q. (2016) Sperm tsRNAs contribute to intergenerational inheritance of an acquired metabolic disorder, Science. 351, 397–400.

10. Sharma, U., Conine, C. C., Shea, J. M., Boskovic, A., Derr, A. G., Bing, X. Y., Belleannee, C., Kucukural, A., Serra, R. W., Sun, F., Song, L., Carone, B. R., Ricci, E. P., Li, X. Z., Fauquier, L., Moore, M. J., Sullivan, R., Mello, C. C., Garber, M. & Rando, O. J. (2016) Biogenesis and function of tRNA fragments during sperm maturation and fertilization in mammals, Science. 351, 391–396.

11. Zhang, Y. & Chen, Q. (2020) Human sperm RNA code senses dietary sugar, Nat Rev Endocrinol. 16, 200–201.

12. Saikia, M., Jobava, R., Parisien, M., Putnam, A., Krokowski, D., Gao, X. H., Guan, B. J., Yuan, Y., Jankowsky, E., Feng, Z., Hu, G. F., Pusztai-Carey, M., Gorla, M., Sepuri, N. B., Pan, T. & Hatzoglou, M. (2014) Angiogenin-cleaved tRNA halves interact with cytochrome c, protecting cells from apoptosis during osmotic stress, Mol Cell Biol. 34, 2450–63.

13. Chen, Q., Zhang, X., Shi, J., Yan, M. & Zhou, T. (2021) Origins and evolving functionalities of tRNA-derived small RNAs, Trends Biochem Sci. 46, 790–804.

14. Li, S., Xu, Z. & Sheng, J. (2018) tRNA-Derived Small RNA: A Novel Regulatory Small Non-Coding RNA, Genes (Basel). 9.

15. Wei, H., Zhou, B., Zhang, F., Tu, Y., Hu, Y., Zhang, B. & Zhai, Q. (2013) Profiling and identification of small rDNA-derived RNAs and their potential biological functions, PLoS One. 8, e56842.

16. Chu, C., Yu, L., Wu, B., Ma, L., Gou, L. T., He, M., Guo, Y., Li, Z. T., Gao, W., Shi, H., Liu, M. F., Wang, H., Chen, C. D., Drevet, J. R., Zhou, Y. & Zhang, Y. (2017) A sequence of 28S rRNA-derived small RNAs is enriched in mature sperm and various somatic tissues and possibly associates with inflammation, J Mol Cell Biol. 9, 256–259.

17. Cherlin, T., Magee, R., Jing, Y., Pliatsika, V., Loher, P. & Rigoutsos, I. (2020) Ribosomal RNA fragmentation into short RNAs (rRFs) is modulated in a sex- and population of origin-specific manner, BMC Biol. 18, 38.

18. Lee, H. C., Chang, S. S., Choudhary, S., Aalto, A. P., Maiti, M., Bamford, D. H. & Liu, Y. (2009) qiRNA is a new type of small interfering RNA induced by DNA damage, Nature. 459, 274–7.

19. Li, S. (2019) Human 28s rRNA 5’ terminal derived small RNA inhibits ribosomal protein mRNA levels, bioRxiv, 618520.

20. Aparicio-Puerta, E., Lebron, R., Rueda, A., Gomez-Martin, C., Giannoukakos, S., Jaspez, D., Medina, J. M., Zubkovic, A., Jurak, I., Fromm, B., Marchal, J. A., Oliver, J. & Hackenberg, M. (2019) sRNAbench and sRNAtoolbox 2019: intuitive fast small RNA profiling and differential expression, Nucleic Acids Res. 47, W530–W535.

21. Schimmel, P. (2018) The emerging complexity of the tRNA world: mammalian tRNAs beyond protein synthesis, Nat Rev Mol Cell Biol. 19, 45–58.

22. Sergiev, P. V., Aleksashin, N. A., Chugunova, A. A., Polikanov, Y. S. & Dontsova, O. A. (2018) Structural and evolutionary insights into ribosomal RNA methylation, Nat Chem Biol. 14, 226–235.

23. Sharma, U., Sun, F., Conine, C. C., Reichholf, B., Kukreja, S., Herzog, V. A., Ameres, S. L. & Rando, O. J. (2018) Small RNAs Are Trafficked from the Epididymis to Developing Mammalian Sperm, Dev Cell. 46, 481–494 e6.

24. Armache, J. P., Jarasch, A., Anger, A. M., Villa, E., Becker, T., Bhushan, S., Jossinet, F., Habeck, M., Dindar, G., Franckenberg, S., Marquez, V., Mielke, T., Thomm, M., Berninghausen, O., Beatrix, B., Soding, J., Westhof, E., Wilson, D. N. & Beckmann, R. (2010) Cryo-EM structure and rRNA model of a translating eukaryotic 80S ribosome at 5.5-A resolution, Proc Natl Acad Sci U S A. 107, 19748–53.

25. Ninomiya, S., Kawano, M., Abe, T., Ishikawa, T., Takahashi, M., Tamura, M., Takahashi, Y. & Nashimoto, M. (2015) Potential small guide RNAs for tRNase ZL from human plasma, peripheral blood mononuclear cells, and cultured cell lines, PLoS One. 10, e0118631.

26. Reilly, J. N., McLaughlin, E. A., Stanger, S. J., Anderson, A. L., Hutcheon, K., Church, K., Mihalas, B. P., Tyagi, S., Holt, J. E., Eamens, A. L. & Nixon, B. (2016) Characterisation of mouse epididymosomes reveals a complex profile of microRNAs and a potential mechanism for modification of the sperm epigenome, Sci Rep. 6, 31794.

27. Bahat, A., Caplan, S. R. & Eisenbach, M. (2012) Thermotaxis of human sperm cells in extraordinarily shallow temperature gradients over a wide range, PLoS One. 7, e41915.

28. Bahat, A., Tur-Kaspa, I., Gakamsky, A., Giojalas, L. C., Breitbart, H. & Eisenbach, M. (2003) Thermotaxis of mammalian sperm cells: a potential navigation mechanism in the female genital tract, Nat Med. 9, 149–50.

29. Bahat, A. & Eisenbach, M. (2006) Sperm thermotaxis, Mol Cell Endocrinol. 252, 115–9.

