## Supplementary figures and images for "The expression and function of 28s5-rtsRNA: an update"

### Supplementary Figure 1

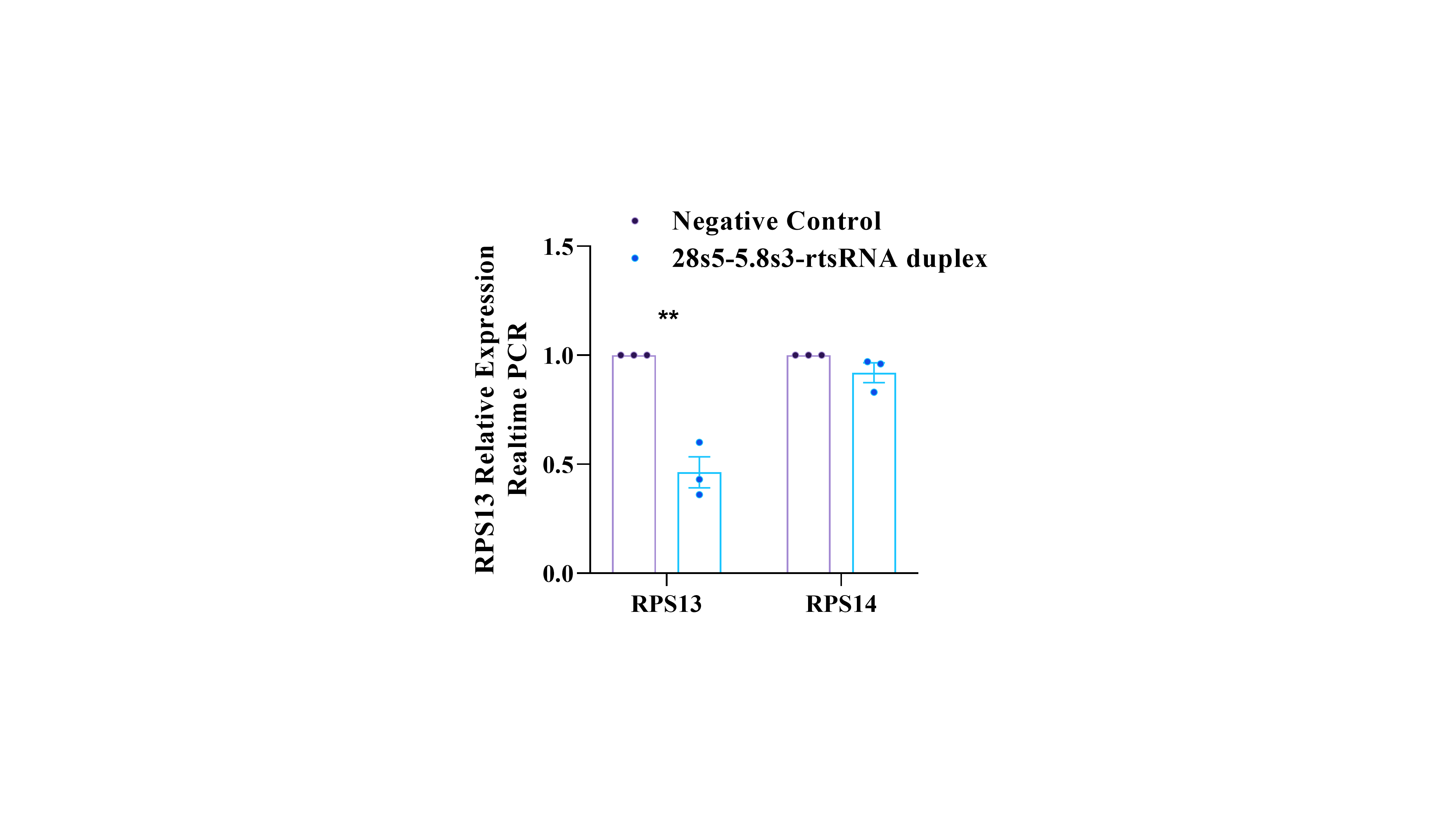

### Supplementary Figure 2

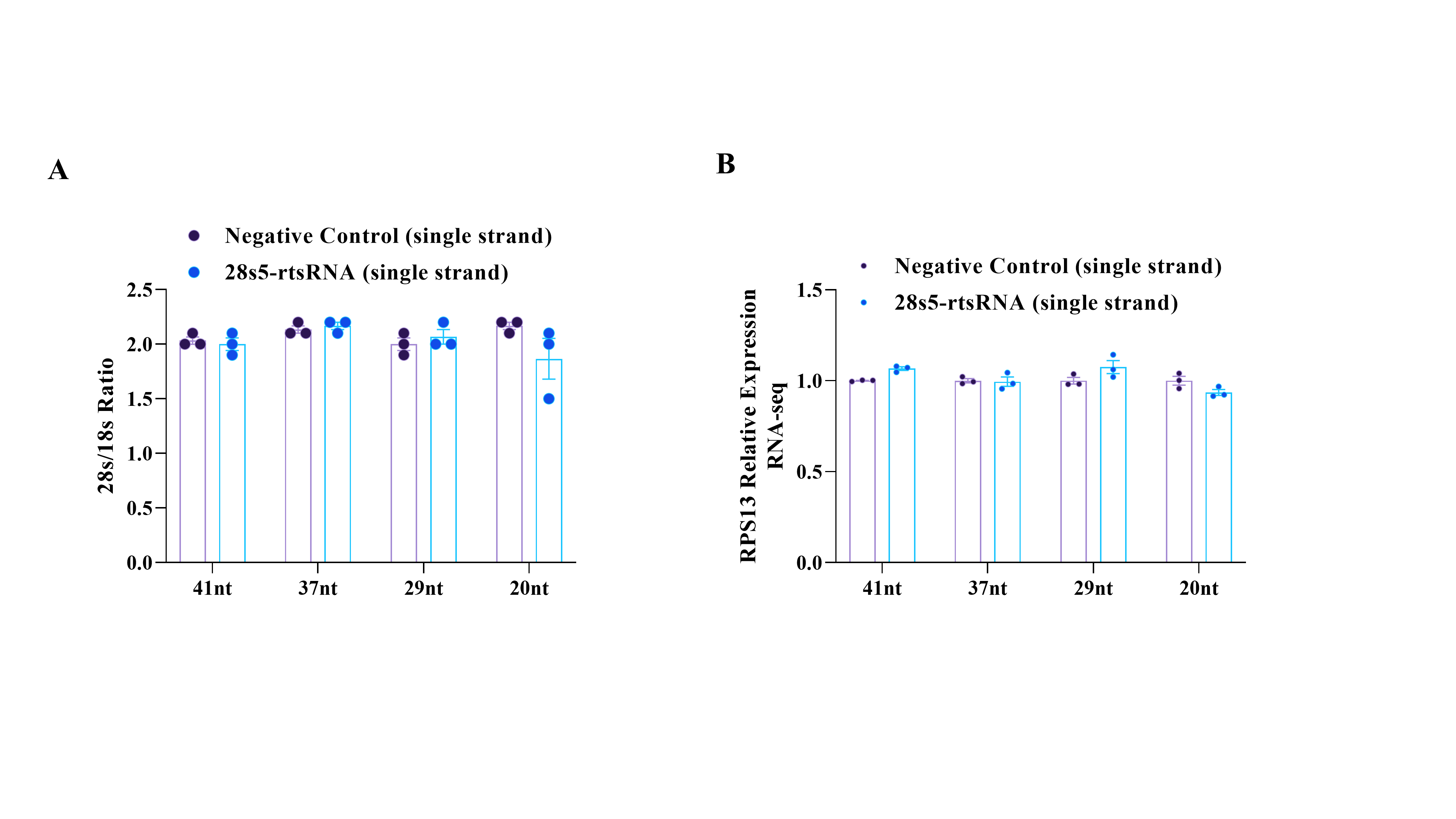

### Supplementary Figure 3

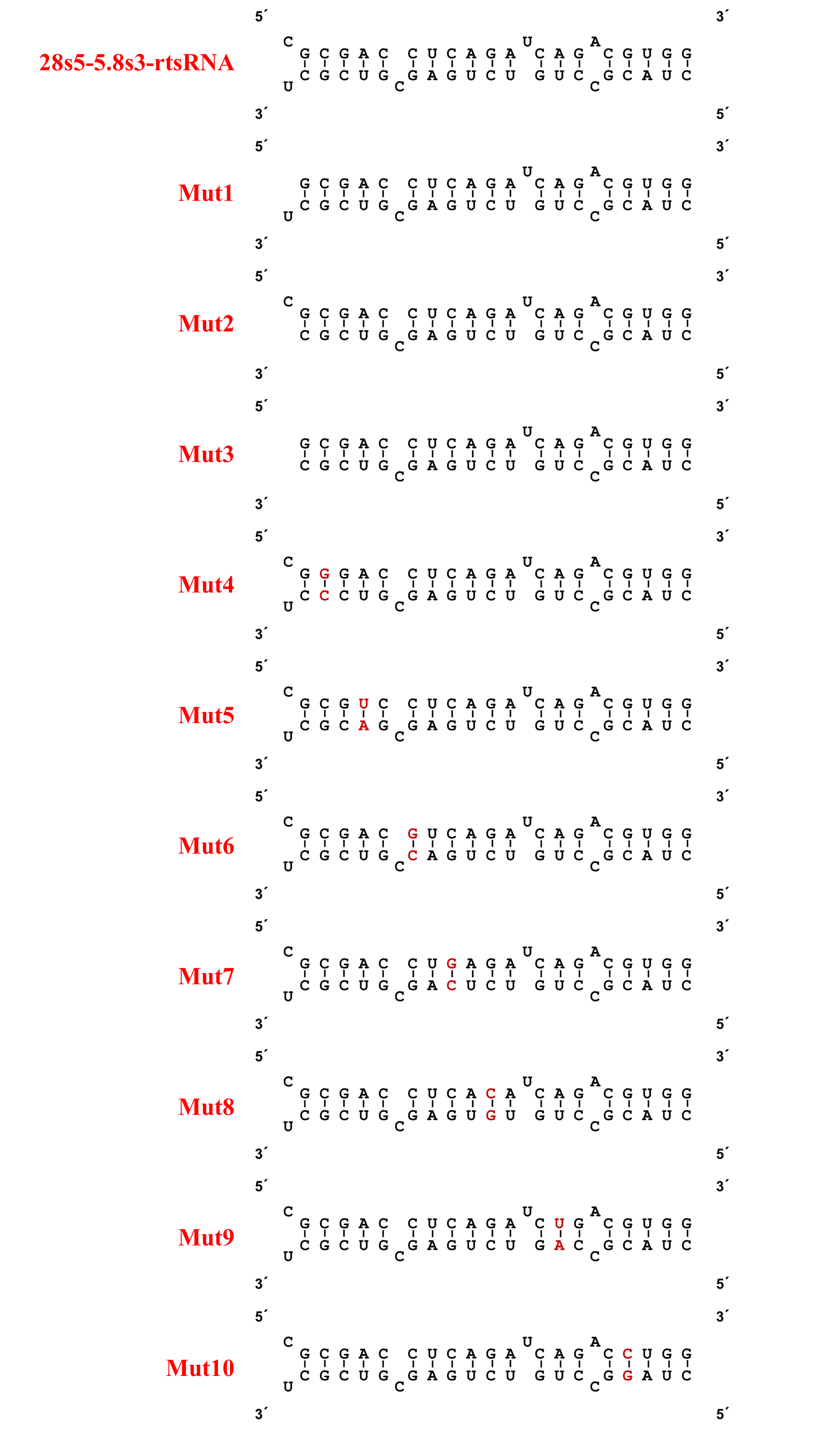

### Supplementary Figure 4

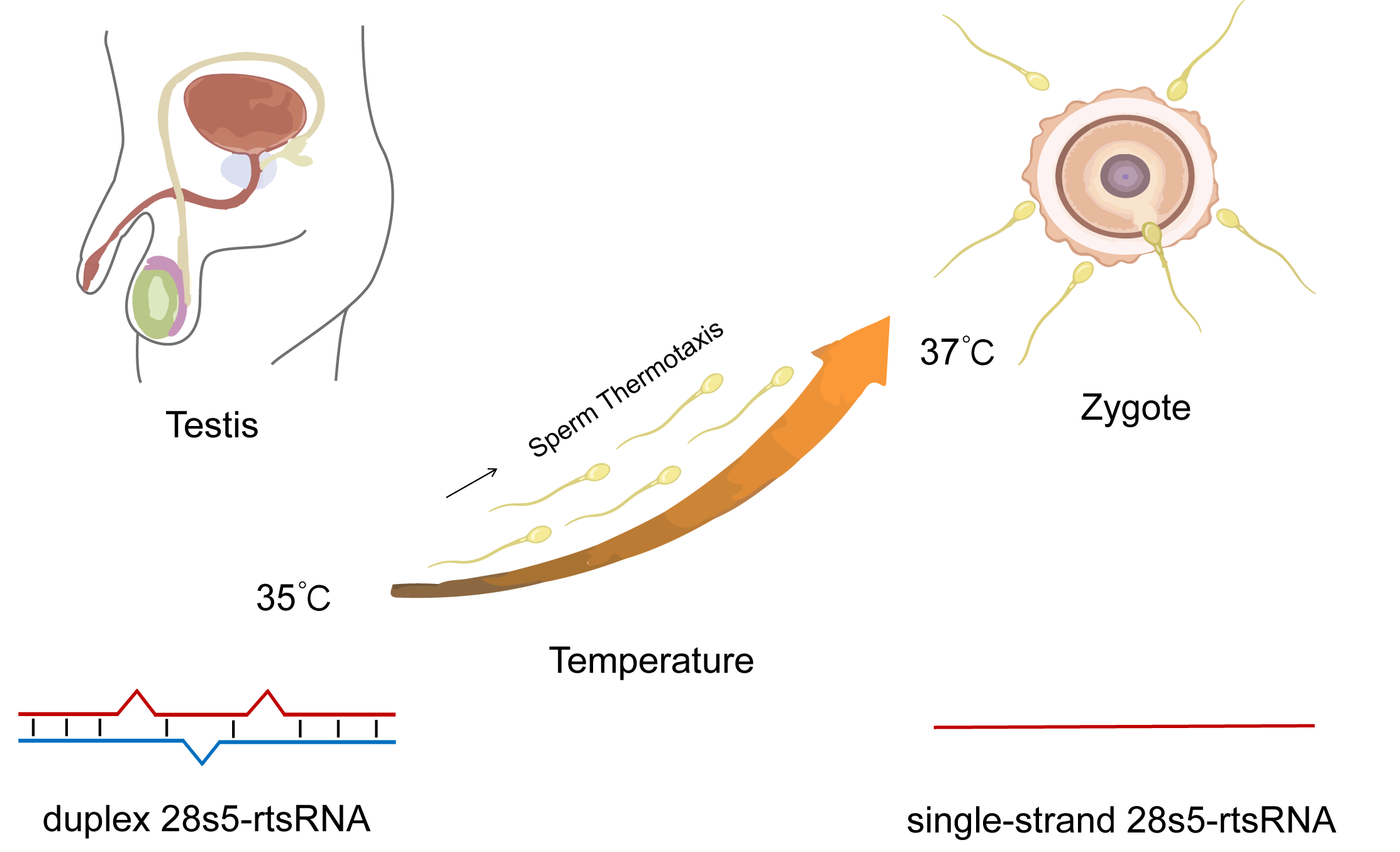

### Supplementary Figure 5

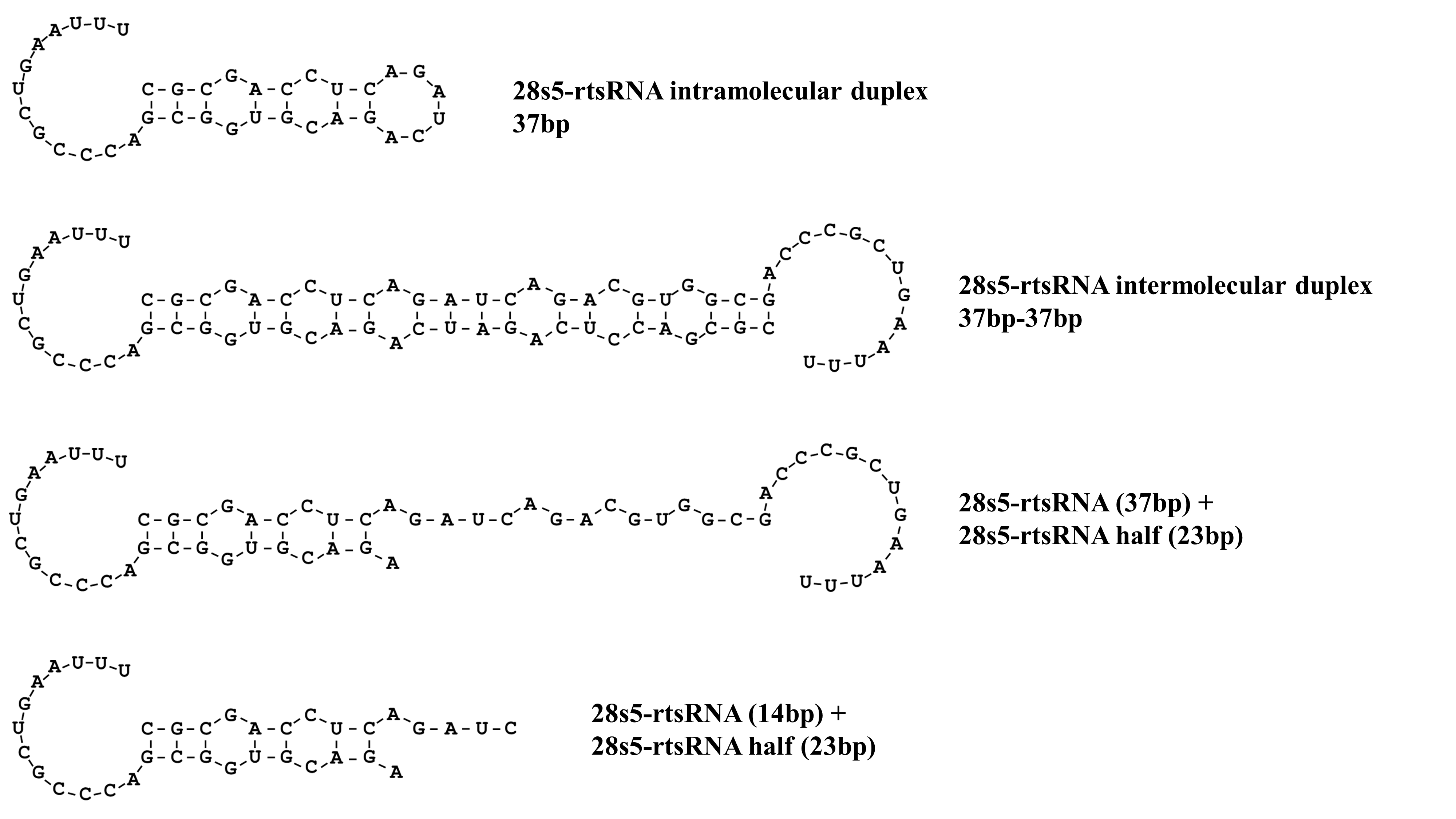
