## Supplementary material for "The expression and function of 28s5-rtsRNA: an update": Table 1

Table 1. Top 10 Small RNAs in Mouse Brain (SRR11004020)

| No | Sequence | Length | Reads | 28s5-rtsRNA |
| --- | --- | --- | --- | --- |
| 1 | CGCGACCUCAGAUCAGACGUGGCGACCCGCUGAAUUU | 37 | 996,414 | ✓ |
| 2 | CGCGACCUCAGAUCAGACGUGGCGACCCGCUGAAUU | 36 | 297,201 | ✓ |
| 3 | CGCGACCUCAGAUCAGACGUGGCGACCCGCUGAAU | 35 | 296,004 | ✓ |
| 4 | UCCCUGGUGGUCUAGUGGUUAGGAUUCGGCGCUCUC | 36 | 227,695 |  |
| 5 | CGCGACCUCAGAUCAGACGUGGCGACCCGCUGAA | 34 | 164,064 | ✓ |
| 6 | UCGGAUCCGUCUGAGCUUGGCUU | 23 | 136,758 |  |
| 7 | UCCCUGGUGGUCUAGUGGUUAGGAUUCGGCGCUC | 34 | 117,205 |  |
| 8 | UGAGGUAGUAGGUUGUAUGGUU | 22 | 99,877 |  |
| 9 | CGCGACCUCAGAUCAGACGUGGCGACCCGCUGAAUUUA | 38 | 97,116 | ✓ |
| 10 | UCCCUGUGGUCUAGUGGUUAGGAUUCGGCGCUCUC | 35 | 90,095 |  |
