## Supplementary material for "The expression and function of 28s5-rtsRNA: an update": Table 2

Table 2. Top 10 Small RNAs in Mouse Mature Sperm (SRR11004052)

| No | Sequence | Length | Reads | 28s5-rtsRNA | 28s5-rtsRNA half |
| --- | --- | --- | --- | --- | --- |
| 1 | CGCGACCUCAGAUCAGACGUGGCGACCCGCUGAA | 34 | 1,317,116 | ✓ |  |
| 2 | AGACGUGGCGACCCGCUGAA | 20 | 834,475 |  | ✓ |
| 3 | AGACGUGGCGACCCGCUGAAUUU | 23 | 609,554 |  | ✓ |
| 4 | AGACGUGGCGACCCGCUGAAUUUAAGC | 27 | 470,011 |  | ✓ |
| 5 | CGCGACCUCAGAUCAGACGUGGCGACCCGCUGAAUUU | 37 | 408,998 | ✓ |  |
| 6 | AGACGUGGCGACCCGCUGAAU | 21 | 402,846 |  | ✓ |
| 7 | ACGUUCGUGUGGAACCUGGCGCUAAACC | 28 | 369,485 |  |  |
| 8 | CGCGACCUCAGAUCAGACGUGGCGACCCGCUGAAU | 35 | 362,090 | ✓ |  |
| 9 | UCGAUGUCGGCUCUUCCUAUC | 21 | 231,950 |  |  |
| 10 | AAGAACGAAAGUCGGAGGUUCGAAGACGAUC | 31 | 198,924 |  |  |
