## Supplementary material for "The expression and function of 28s5-rtsRNA: an update": Table 3

Table 3. KEGG Pathway Analysis of 28s5-5.8s3-rtsRNA Downregulated Genes

| NO | ID | Name | pValue | FDR B&H | FDR B&Y | Genes from Input | Genes in Annotation |
| --- | --- | --- | --- | --- | --- | --- | --- |
| 1 | 99052 | Lysosome | 2.06E-13 | 3.34E-11 | 2.82E-10 | 30 | 123 |
| 2 | 83036 | Ribosome | 3.82E-10 | 4.50E-08 | 3.80E-07 | 29 | 153 |
| 3 | 82973 | Glutathione metabolism | 1.71E-06 | 1.23E-04 | 1.04E-03 | 13 | 54 |
| 4 | 83055 | p53 signaling pathway | 2.99E-05 | 1.85E-03 | 1.56E-02 | 13 | 69 |
| 5 | 413377 | Heparan sulfate degradation | 3.21E-05 | 1.94E-03 | 1.63E-02 | 5 | 9 |
| 6 | 213306 | Measles | 3.47E-05 | 2.04E-03 | 1.73E-02 | 19 | 134 |
| 7 | 658418 | Viral carcinogenesis | 6.20E-05 | 3.49E-03 | 2.95E-02 | 24 | 201 |
| 8 | 82981 | Glycosaminoglycan degradation | 2.31E-04 | 1.20E-02 | 1.01E-01 | 6 | 19 |
| 9 | 83117 | Acute myeloid leukemia | 3.34E-04 | 1.60E-02 | 1.35E-01 | 10 | 55 |
| 10 | 694606 | Hepatitis B | 8.07E-04 | 3.22E-02 | 2.72E-01 | 17 | 144 |
