## Supplementary material for "The expression and function of 28s5-rtsRNA: an update": Table 4

Table 4. KEGG Pathway Analysis of 28s5-5.8s3-rtsRNA Upregulated Genes

| NO | ID | Name | pValue | FDR B&H | FDR B&Y | Genes from Input | Genes in Annotation |
| --- | --- | --- | --- | --- | --- | --- | --- |
| 1 | 83054 | Cell cycle | 8.93E-07 | 1.18E-04 | 1.00E-03 | 24 | 124 |
| 2 | 177876 | RNA transport | 3.78E-06 | 3.88E-04 | 3.30E-03 | 28 | 171 |
| 3 | 199378 | Ribosome biogenesis in eukaryotes | 2.85E-05 | 2.03E-03 | 1.72E-02 | 19 | 104 |
| 4 | 119304 | Progesterone-mediated oocyte maturation | 3.20E-05 | 2.22E-03 | 1.89E-02 | 18 | 96 |
| 5 | 126909 | Oocyte meiosis | 3.61E-05 | 2.45E-03 | 2.08E-02 | 21 | 124 |
| 6 | 83103 | Pathogenic Escherichia coli infection | 1.48E-04 | 7.19E-03 | 6.11E-02 | 12 | 55 |
| 7 | 373901 | HTLV-I infection | 4.52E-04 | 1.53E-02 | 1.30E-01 | 31 | 256 |
| 8 | 83118 | Small cell lung cancer | 8.24E-04 | 2.34E-02 | 1.99E-01 | 14 | 84 |
| 9 | 585562 | Epstein-Barr virus infection | 1.06E-03 | 2.86E-02 | 2.43E-01 | 25 | 201 |
| 10 | 1272486 | Insulin resistance | 1.22E-03 | 3.04E-02 | 2.58E-01 | 16 | 107 |
