## Supplementary Table 1 for "The expression and function of 28s5-rtsRNA: an update"

Supplementary Table 1. High throughput sequencing data used in this study

| Species | Tissue | SRA | Referred in text | Reference |
| --- | --- | --- | --- | --- |
| Mouse | Brain | SRR11004020 | Table 1 | 2 |
| Mouse | Mature Sperm | SRR11004052 | Table 2 | 2 |
| Mouse | Brain traditional rep 1 | SRR11004011 | Figure 1, 3 | 2 |
| Mouse | Brain traditional rep 2 | SRR11004012 | Figure 1, 3 | 2 |
| Mouse | Brain traditional rep 3 | SRR11004013 | Figure 1, 3 | 2 |
| Mouse | Brain +AlkB rep 1 | SRR11004014 | Figure 1, 3 | 2 |
| Mouse | Brain +AlkB rep 2 | SRR11004015 | Figure 1, 3 | 2 |
| Mouse | Brain +AlkB rep 3 | SRR11004016 | Figure 1, 3 | 2 |
| Mouse | Brain +T4PNK rep 1 | SRR11004017 | Figure 1, 3 | 2 |
| Mouse | Brain +T4PNK rep 2 | SRR11004018 | Figure 1, 3 | 2 |
| Mouse | Brain +T4PNK rep 3 | SRR11004019 | Figure 1, 3 | 2 |
| Mouse | Brain PANDORA-seq rep 1 | SRR11004020 | Figure 1, 3 | 2 |
| Mouse | Brain PANDORA-seq rep 2 | SRR11004021 | Figure 1, 3 | 2 |
| Mouse | Brain PANDORA-seq rep 3 | SRR11004022 | Figure 1, 3 | 2 |
| Mouse | Liver traditional rep 1 | SRR11004023 | Figure 1, 3 | 2 |
| Mouse | Liver traditional rep 2 | SRR11004024 | Figure 1, 3 | 2 |
| Mouse | Liver traditional rep 3 | SRR11004025 | Figure 1, 3 | 2 |
| Mouse | Liver +AlkB rep 1 | SRR11004026 | Figure 1, 3 | 2 |
| Mouse | Liver +AlkB rep 2 | SRR11004027 | Figure 1, 3 | 2 |
| Mouse | Liver +AlkB rep 3 | SRR11004028 | Figure 1, 3 | 2 |
| Mouse | Liver +T4PNK rep 1 | SRR11004029 | Figure 1, 3 | 2 |
| Mouse | Liver +T4PNK rep 2 | SRR11004030 | Figure 1, 3 | 2 |
| Mouse | Liver +T4PNK rep 3 | SRR11004031 | Figure 1, 3 | 2 |
| Mouse | Liver PANDORA-seq rep 1 | SRR11004032 | Figure 1, 3  Supplementary Table 5 | 2 |
| Mouse | Liver PANDORA-seq rep 2 | SRR11004033 | Figure 1, 3 | 2 |
| Mouse | Liver PANDORA-seq rep 3 | SRR11004034 | Figure 1, 3 | 2 |
| Mouse | Mature sperm traditional rep 1 | SRR11004043 | Figure 1, 3 | 2 |
| Mouse | Mature sperm traditional rep 2 | SRR11004044 | Figure 1, 3 | 2 |
| Mouse | Mature sperm traditional rep 3 | SRR11004045 | Figure 1, 3 | 2 |
| Mouse | Mature sperm +AlkB rep 1 | SRR11004046 | Figure 1, 3 | 2 |
| Mouse | Mature sperm +AlkB rep 2 | SRR11004047 | Figure 1, 3 | 2 |
| Mouse | Mature sperm +AlkB rep 3 | SRR11004048 | Figure 1, 3 | 2 |
| Mouse | Mature sperm +T4PNK rep 1 | SRR11004049 | Figure 1, 3 | 2 |
| Mouse | Mature sperm +T4PNK rep 2 | SRR11004050 | Figure 1, 3 | 2 |
| Mouse | Mature sperm +T4PNK rep 3 | SRR11004051 | Figure 1, 3 | 2 |
| Mouse | Mature sperm PANDORA-seq rep 1 | SRR11004052 | Figure 1, 3 | 2 |
| Mouse | Mature sperm PANDORA-seq rep 2 | SRR11004053 | Figure 1, 3 | 2 |
| Mouse | Mature sperm PANDORA-seq rep 3 | SRR11004054 | Figure 1, 3 | 2 |
| Mouse | ESC traditional rep 1 | SRR11004091 | Figure 1, 3 | 2 |
| Mouse | ESC traditional rep 2 | SRR11004092 | Figure 1, 3 | 2 |
| Mouse | ESC traditional rep 3 | SRR11004093 | Figure 1, 3 | 2 |
| Mouse | ESC +AlkB rep 1 | SRR11004094 | Figure 1, 3 | 2 |
| Mouse | ESC +AlkB rep 2 | SRR11004095 | Figure 1, 3 | 2 |
| Mouse | ESC +AlkB rep 3 | SRR11004096 | Figure 1, 3 | 2 |
| Mouse | ESC +T4PNK rep 1 | SRR11004097 | Figure 1, 3 | 2 |
| Mouse | ESC +T4PNK rep 2 | SRR11004098 | Figure 1, 3 | 2 |
| Mouse | ESC +T4PNK rep 3 | SRR11004099 | Figure 1, 3 | 2 |
| Mouse | ESC PANDORA-seq rep 1 | SRR11004100 | Figure 1, 3 | 2 |
| Mouse | ESC PANDORA-seq rep 2 | SRR11004101 | Figure 1, 3 | 2 |
| Mouse | ESC PANDORA-seq rep 3 | SRR11004102 | Figure 1, 3 | 2 |
| Mouse | Cauda Epididymis Sperm -PNK | SRR6984142 | Supplementary Table 3 | 22 |
| Mouse | Cauda Epididymis Sperm +PNK | SRR6984143 | Supplementary Table 4 | 22 |
| Human | ESC | SRR11004132 | Supplementary Table 6 | 2 |
