## Supplementary Table 2 for "The expression and function of 28s5-rtsRNA: an update"

Supplementary Table 2. RNA sequences used in this study

| ID | Format | Sequence | Passenger |
| --- | --- | --- | --- |
| 28s5-5.8s3-rtsRNA | duplex | CGCGACCUCAGAUCAGACGUGG | CUACGCCUGUCUGAGCGUCGCU |
| Mut1 | duplex | GCGACCUCAGAUCAGACGUGG | CUACGCCUGUCUGAGCGUCGCU |
| Mut2 | duplex | CGCGACCUCAGAUCAGACGUGG | CUACGCCUGUCUGAGCGUCGC |
| Mut3 | duplex | GCGACCUCAGAUCAGACGUGG | CUACGCCUGUCUGAGCGUCGC |
| Mut4 | duplex | CGgGACCUCAGAUCAGACGUGG | CUACGCCUGUCUGAGCGUCcCU |
| Mut5 | duplex | CGCGuCCUCAGAUCAGACGUGG | CUACGCCUGUCUGAGCGaCGCU |
| Mut6 | duplex | CGCGACgUCAGAUCAGACGUGG | CUACGCCUGUCUGAcCGUCGCU |
| Mut7 | duplex | CGCGACCUgAGAUCAGACGUGG | CUACGCCUGUCUcAGCGUCGCU |
| Mut8 | duplex | CGCGACCUCAcAUCAGACGUGG | CUACGCCUGUgUGAGCGUCGCU |
| Mut9 | duplex | CGCGACCUCAGAUCuGACGUGG | CUACGCCaGUCUGAGCGUCGCU |
| Mut10 | duplex | CGCGACCUCAGAUCAGACcUGG | CUAgGCCUGUCUGAGCGUCGCU |
| Negative Control | duplex | UUCUCCGAACGUGUCACGUTT | ACGUGACACGUUCGGAGAATT |
| siLuc | duplex | CUUACGCUGAGUACUUCGAUU | UCGAAGUACUCAGCGUAAGUU |
| NC-41nt | single | AUUCGGAGCGCCGAGUCGUACUGCCGGACACACGCAACUAU | |
| 28s5-rtsRNA-41nt | single | CGCGACCUCAGAUCAGACGUGGCGACCCGCUGAAUUUAAGC | |
| NC-37nt | single | GUUCGUUCACACGAGACGCCUAUCGGCACGUACGACG | |
| 28s5-rtsRNA-37nt | single | CGCGACCUCAGAUCAGACGUGGCGACCCGCUGAAUUU | |
| NC-29nt | single | GCCACGGAUCCGCACCUGCGACGCGAGUA | |
| 28s5-rtsRNA-29nt | single | CGCGACCUCAGAUCAGACGUGGCGACCCG | |
| NC-20nt | single | UUGUACUACACAAAAGUACUG | |
| 28s5-rtsRNA-20nt | single | CGCGACCUCAGAUCAGACGU | |
