## Supplementary Table 3 for "The expression and function of 28s5-rtsRNA: an update"

Supplementary Table 3. Top 10 Small RNAs in Cauda Epididymis Sperm -PNK (SRR6984142)

| No | Sequence | Length | Reads | 28s5-rtsRNA |
| --- | --- | --- | --- | --- |
| 1 | GCAUUGGUGGUUCAGUGG | 18 | 539,453 |  |
| 2 | UCCCUGGUGGUCUAGUGGUUAGGAUUCGGC | 30 | 194,324 |  |
| 3 | GCAUUGUGGUUCAGUGG | 17 | 190,179 |  |
| 4 | UCCCUGUGGUCUAGUGGUUAGGAUUCGGC | 29 | 113,062 |  |
| 5 | GUUUCCGUAGUGUAGUGGUUAUCACGUUCGCCU | 33 | 94,052 |  |
| 6 | UCCCUGGUGGUCUAGUGGUUAGGAUUCGGCG | 31 | 79,918 |  |
| 7 | GUUUCCGUAGUGUAGUGGUUAUCACGUUCGCC | 32 | 51,640 |  |
| 8 | UCCCUGUGGUCUAGUGGUUAGGAUUCGGCG | 30 | 46,327 |  |
| 9 | UACCCUGUAGAACCGAAUUUGU | 22 | 41,488 |  |
| 10 | GUUUCCGUAGUGUAGUGGUUAUCACGUUCGCCUC | 34 | 40,007 |  |
