## Supplementary Table 4 for "The expression and function of 28s5-rtsRNA: an update"

Supplementary Table 4. Top 10 Small RNAs in Cauda Epididymis Sperm +PNK (SRR6984143)

| No | Sequence | Length | Reads | 28s5-rtsRNA |
| --- | --- | --- | --- | --- |
| 1 | CGCGACCUCAGAUCAGACGUGGCGACCCGCUGAAUUUAAGC | 41 | 871,424 | ✓ |
| 2 | CGCGACCUCAGAUCAGACGUGGCGACCCGCUGAAUUU | 37 | 174,873 | ✓ |
| 3 | GCAUUGGUGGUUCAGUGG | 18 | 73,257 |  |
| 4 | GUUUCCGUAGUGUAGUGGUUAUCACGUUCGCCUC | 34 | 63,549 |  |
| 5 | GCCCGGCUAGCUCAGUCGGUAGAGCAUGAGACUC | 34 | 37,611 |  |
| 6 | GCAUUGUGGUUCAGUGG | 17 | 27,730 |  |
| 7 | CGCGACCUCAGAUCAGACGUGGCGACCCGCUGAAUU | 36 | 25,539 | ✓ |
| 8 | GUUUCCGUAGUGUAGUGGUUAUCACGUUCGCC | 32 | 24,815 |  |
| 9 | GUUUCCGUAGUGUAGUGGUUAUCACGUUCGCCUGAC | 36 | 24,136 |  |
| 10 | GUUUCCGUAGUGUAGUGGUUAUCACGUUCGCCU | 33 | 19,131 |  |
