## Supplementary Table 5 for "The expression and function of 28s5-rtsRNA: an update"

Supplementary Table 5. Top 10 Small RNAs in Mouse Liver (SRR11004032)

| No | Sequence | Length | Reads | 28s5-rtsRNA |
| --- | --- | --- | --- | --- |
| 1 | CGCGACCUCAGAUCAGACGUGGCGACCCGCUGAAUUU | 37 | 4,431,146 | ✓ |
| 2 | AGCCGACUUAGAACUGGUGCGGACCAGGGGAAUCCGACUGUUU | 43 | 1,846,511 |  |
| 3 | UGGAGUGUGACAAUGGUGUUUGU | 23 | 1,295,899 |  |
| 4 | UGGAGUGUGACAAUGGUGUUUG | 22 | 1,283,357 |  |
| 5 | UGGAGUGUGACAAUGGUGUUUGA | 23 | 980,976 |  |
| 6 | UGGAGUGUGACAAUGGUGUUU | 21 | 891,413 |  |
| 7 | GUGCGUCGAUGAAGAACGCAGCUAGCUGCGAGAAUU | 36 | 254,804 |  |
| 8 | ACUUGGAUGGGAGACCGCCUGGGAAUACCGGGUGCUGUAGGCUU | 44 | 228,534 |  |
| 9 | AGACGUGGCGACCCGCUGAAUUUAAGCAUAUU | 32 | 202,886 |  |
| 10 | UCAGUGCACUACAGAACUUUGU | 22 | 192,652 |  |
