## Supplementary Table 6 for "The expression and function of 28s5-rtsRNA: an update"

Supplementary Table 6. Top 10 Small RNAs in Human ESC (SRR11004132)

| No | Sequence | Length | Reads | 28s5-rtsRNA |
| --- | --- | --- | --- | --- |
| 1 | CGCGACCUCAGAUCAGACGUGGCGACCCGCUGAAUUU | 37 | 1,654,674 | ✓ |
| 2 | CGCGACCUCAGAUCAGACGUGGCGACCCGCUGAA | 34 | 569,867 | ✓ |
| 3 | AGCCGACUUAGAACUGGUGCGGACCAGGGGAAUCCGACUGUUU | 43 | 497,593 |  |
| 4 | CGCGCCGGGCCCGGGUCUUCCCG | 23 | 423,210 |  |
| 5 | CGCGACCUCAGAUCAGACGUGGCGACCCGCUGAAU | 35 | 321,465 | ✓ |
| 6 | AGGGGGCGGGCUCCGGCGGG | 20 | 313,290 |  |
| 7 | CGCGACCUCAGAUCAGACGUGGCGACCCGCUGAAUU | 36 | 270,236 | ✓ |
| 8 | GGGGGCGGGCUCCGGCGGG | 19 | 195,215 |  |
| 9 | UCGGGCUGGGGCGCGAAGCGGGGC | 24 | 172,283 |  |
| 10 | GCGACCUCAGAUCAGACGUGGCGACCCGCUGAA | 33 | 163,578 |  |
